# Application of machine learning methods in clinical trials for precision medicine

**DOI:** 10.1101/2021.10.06.463354

**Authors:** Yizhuo Wang, Bing Z. Carter, Ziyi Li, Xuelin Huang

**Affiliations:** Department of Biostatistics, The University of Texas MD Anderson Cancer Center, Houston, TX 77030, USA; Department of Leukemia, The University of Texas MD Anderson Cancer Center, Houston, TX 77030, USA

## Abstract

**Objective:** A key component for precision medicine is a good prediction algorithm for patients’ response to treatments. We aim to implement machine learning (ML) algorithms into the response-adaptive randomization (RAR) design and improve the treatment outcomes.

**Materials and Methods:** We incorporated nine ML algorithms to model the relationship of patient responses and biomarkers in clinical trial design. Such a model predicted the response rate of each treatment for each new patient and provide guidance for treatment assignment. Realizing that no single method may fit all trials well, we also built an ensemble of these nine methods. We evaluated their performance through quantifying the benefits for trial participants, such as the overall response rate and the percentage of patients who receive their optimal treatments.

**Results:** Simulation studies showed that the adoption of ML methods resulted in more personalized optimal treatment assignments and higher overall response rates among trial participants. Compared with each individual ML method, the ensemble approach achieved the highest response rate and assigned the largest percentage of patients to their optimal treatments. For the real-world study, we successfully showed the potential improvements if the proposed design had been implemented in the study.

**Conclusion:** In summary, the ML-based RAR design is a promising approach for assigning more patients to their personalized effective treatments, which makes the clinical trial more ethical and appealing. These features are especially desirable for late-stage cancer patients who have failed all the FDA-approved treatment options and only can get new treatments through clinical trials.

## 1 INTRODUCTION

It is known that patients respond differently to the same treatments [2]. The demand for selecting the optimal treatment for each and every patient has resulted in a rapidly developing field called precision medicine, also known as personalized medicine [18]. This field aims to provide guidance to select the most effective treatment based on distinctive patient biomarkers. As clinical trials also evolve in the age of precision medicine, there is a substantial need for novel trial designs to deliver more ethical and precise care. Compared with classical non-adaptive trials, adaptive trials have become popular among clinicians as they integrate accumulating patient data to modify the parameters of the trial protocol, provide personalized treatment assignment, and ultimately optimize patients’ outcomes. For example, the adaptive designs in phase 2/3 clinical trials take advantage of the interim treatment response data during the course of the trial and allocate more patients to the presumably more effective treatments [35].

Among different adaptive designs, one common adaptation is response-adaptive randomization (RAR). It refers to the adjustments of treatment allocations based on intermediate patient responses and new patients’ characteristics collected during the clinical trials. This RAR design is useful when the interaction between biomarkers and treatments are only putative or not known at the beginning of a trial, and it is also practical when there are multiple treatments to be considered. Its ultimate objective is to provide more patients with their personalized optimal therapies according to their biomarker profiles. The starting point of RAR can be traced back to Thompson (1933), who proposed employing a posterior probability estimated from the interim data to assign patients to the more effective treatment [49]. Following his idea, the application of Bayesian methods with an inherent adaptive nature has boomed in area of RAR designs [5, 60, 57, 53, 7, 30, 28, 48, 11].

Currently, there are several major successes in applying Bayesian RAR concepts in clinical trials, from protocol development through legitimate registration. The BATTLE-1 trial (Biomarker-integrated Approaches of Targeted Therapy for Lung Cancer Elimination) for patients with advanced non-small cell lung cancer (NSCLC) and the I-SPY 2 trial (Investigation of Serial Studies to Predict Your Therapeutic Response with Imaging and Molecular Analysis) for patients with breast cancer in the setting of neoadjuvant chemotherapy are two biomarker-based, Bayesian RAR clinical trials [26, 4]. However, Bayesian RAR designs have a number of challenges and limitations. Due to the modeling restrictions, Bayesian RAR methods usually consider only a very small number of biomarkers. With complex diseases or symptoms, hundreds or even thousands of biomarkers may need to be considered at the same time for treatment assignment [37]. Also, some Bayesian RAR methods adjust the design by separating the cohort based on the existence of biomarker(s), and thus these methods rely heavily on how well the biomarker(s) interact(s) with the treatments. If the biomarker is chosen incorrectly, it is possible to make wrong adjustments afterwards [29, 2].

As the development of modern sequencing technology, clinicians have faced a massive volume of high dimensional data with a complex, nonlinear structure. How to build an effective and scalable algorithm for randomization becomes a fundamental question for the research of RAR trial designs. Machine learning (ML) methods have been applied to solve many real-world problems and have successfully demonstrated their strengths in processing large data sets, as well as capturing non-linear data structures. With the expectations and resources to analyze this large amount of complex healthcare data, ML methods have established their supremacy in disease prediction [34], disease classification [13], imaging diagnosis [32], drug manufacturing [38], medication assignment [31], and genomic feature identification tasks [59].

Although several supervised ML approaches have been applied to drug response prediction [8, 47, 58, 39, 46, 40, 54], little of the work has explored incorporating ML methods into RAR trial designs. In this study, we implemented nine ML algorithms into RAR designs and further presented an ML-ensemble RAR design combining these nine ML algorithms. Specifically, ML methods help to match patient biomarker profiles with prediction of treatment outcomes and, in turn, have determined treatment allocation for future patients. These ML methods are able to address large data and complex structures. We have successfully demonstrated, in both simulation study and a real-world example, that ML-based RAR designs have higher response rates as there are more patients receiving effective treatments. The ensemble method outperformed all other single ML methods.

## 2 MATERIALS AND METHODS

### 2.1 Adaptive design: response-adaptive randomization

In clinical trial design, adaptive design means making changes to the trial protocol after the trial has started and some data have been collected. These changes are based on the information from the collected data, including (1) the total sample size, (2) interim analyses, (3) patient allocation to different treatment arms, and more [43]. For (3), it refers to the RAR design in which the treatment allocation probability varies in order to favor the treatment estimated to be more effective and to increase the response rate in patients. The initial concept can be traced back to Thompson (1933) [50] and Robbins (1952) [41], and led to others. Some famous RAR trials include the extracorporeal membrane oxygenation (ECMO) trial, which tested the efficacy of ECMO in patients with severe acute respiratory distress syndrome (ARDS) [9], and the first large-scale double-blind, placebo-controlled study which tested the superiority of fluoxetine over placebo in children and adolescents with depression [12]. A general scheme of the RAR designs is shown in Figure 1.

**Figure 1:**
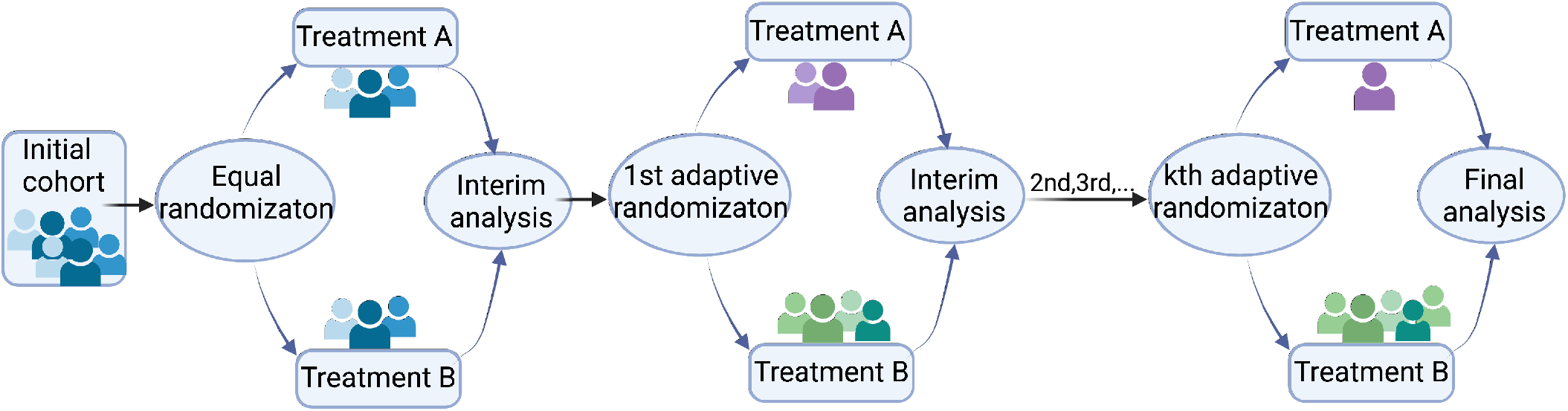
Response-adaptive randomization (RAR) design. The number of adaptive randomization is adjustable per application.

### 2.2 Benchmark design: equal randomization

Randomization as a standard means for addressing the selection bias in treatment assignments has been extensively used in clinical trials [43]. It helps to achieve balance among treatment groups and accounts for the genuine uncertainty about which treatment is better at the beginning of the trial. Randomly assigning patients to treatment arms on a 1:1 basis is known as equal randomization (ER). Friedman, Furberg and DeMets (1981, p.41) presented that equal allocation in principle maximizes statistical power, and is consistent with the concept of equipoise that should exist before the trial starts [16]. Here, we used the ER design as a benchmark randomization design to evaluate the performance of ML-based RAR designs.

### 2.3 Allocation rule

The key of the proposed method is to model the relationship of patient responses and biomarkers. Such a model will then predict the response rate of each treatment for each new patient and provide guidance for treatment assignment. In detail, current enrolled patients’ biomarker profiles and treatment response data were used to train ML models, which later were used to predict future patients’ treatment responses based on their biomarker profiles. Given treatment *A* and *B*, the probability of allocating each treatment for patient *i* is shown as below:

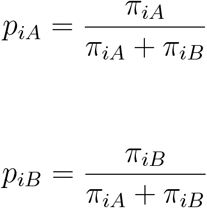

where *π*_*iA*_ and *π*_*iB*_ respectively denote the response probability of treatment *A* and *B* for patient *i* predicted by the ML model.

### 2.4 Machine learning algorithms and a machine learning ensemble

We selected nine mainstream ML algorithms and implemented them in the RAR design to predict treatment response. The prediction models were built using the best parameters for each model, which were obtained by a tenfold cross-validation. These selected ML algorithms can be roughly divided into two categories:

1. Parametric models: logistic regression [10], LASSO regression [51], Ridge regression [23].
2. Non-parametric models: gradient boosting machine (GBM) [15], random forest (RF) [22], support vector machine (SVM) [6], naive bayes [25], k-nearest neighbors (KNN) [1], and artificial neural networks (NN) [21].

For logistic regression, Ridge regression and Lasso regression, they are all considered parametric models. In detail, logistic regression assumes the linearity of independent variables and log odds. It is a particular form of GLM [52]. Ridge regression and LASSO regression assume that there is a linear relationship between the “dependent” variable and the explanatory variables. They are two regularization methods of GLM to prevent an over-fitting issue by adding penalties on the predictor variables that are less significant [23, 51].

KNN, which classifies data points based on the points that are most similar to it, is a typical non-parametric model such that there is no assumption for underlying data distribution, and the number of parameters grows with the size of the data [1]. With NNs, however, there has been some debate regarding whether they belong to parametric or non-parametric methods. NNs typically consist of three layers: input layer, hidden layer, and output layer. Here we classify NNs as a non-parametric method, as the network architecture grows adaptively to match the complexity of given data [21].

Both GBM and RF are nonparametric methods that consist of sets of decision trees. Specifically, GBM builds one tree at a time and each new tree helps to correct errors made by previously trained tree by adding weights to the observations with the worst prediction from the previous iteration; RF trains each tree independently using a random sample of the data, and the results are aggregated in the end [15, 22].

NB and SVM can be either parametric or non-parametric depending on whether they use kernel tricks. For the NB classifier, it becomes non-parametric if using a kernel density estimation (KDE) to obtain a more realistic estimate of the probability of an observation belonging to a class [25]. And for SVM, the basic idea is finding a hyperplane that best divides a dataset into two classes. It is considered a non-parametric when using the kernel trick to find this hyperplane. This is because the kernel is constructed by computing the pair-wise distances between the training points, and the complexity of the model grows with the size of the dataset [6].

Combing these nine models, an ML ensemble method was built and implemented in the RAR design to obtain a better treatment allocation rule. We defined the treatment allocation probability function for patient *i* as follows:

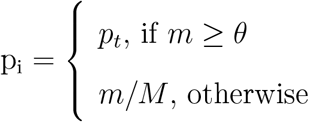

where *m* is the number of agreed models, *M* = 9 is the total number of models, *θ* is the threshold number of agreed models, and *p*_*t*_ is the threshold treatment allocation probability. Here we chose *θ* = 7 and *p*_*t*_ = 0.85; these threshold values can be adjusted accordingly for different application purposes.

Apart from comparing with the ER “benchmark” design, the current study also examined whether the ML ensemble could assign more patients to the best available treatment beyond other ML methods in adaptive design with the same assessment of individuals.

### 2.5 Inverse Probability of Treatment Weighting

Similar to the observational study in which certain outcomes are measured without attempting to change the outcome, the treatment selection of future patients in RAR trials is often influenced by individual characteristics of the initial block of patients [3]. As a result, when estimating the impact of treatment on responses, systemic variations in baseline characteristics between differently-treated individuals must be taken into account. Here we applied the inverse probability of treatment weighted (IPTW) method to decrease or remove the effects of confounding when using the observational data to estimate the treatment effects. The idea of IPTW is to use weights based on the propensity score to create a synthetic sample in which the distribution of baseline characteristics is independent of treatment [42]. The propensity score refers to the probability of treatment allocation tied to the observed individual characteristics. And the weight based on it is defined as follows:

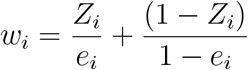

where *Z*_*i*_ is the treatment indicator and *e*_*i*_ is the propensity score for the *i* th subject. Different estimators for treatment effects based on IPTW have been developed; here we used an estimator of the average treatment effect (ATE), which is defined as *E* [*Z*_*i*_(1) − *Z*_*i*_(0)] where *Z*_*i*_(1) − *Z*_*i*_(0) is the effect of treatment.[24] Incorporated with the IPTW idea, the ATE estimator is defined as follows:

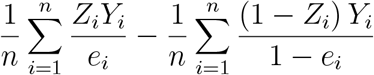

where *Y*_*i*_ denotes the response variable of the *i* th subject, *n* denotes the total number of subjects, and *e*_*i*_ still denotes the propensity score [33].

### 2.6 Evaluation metrics

Two commonly used criterion in the field of precision medicine, namely the overall response rate and the percentage of individuals receiving optimal treatments, are our primary evaluation metrics. The formulas of the response rate and the optimal treatment percentage are as follows:

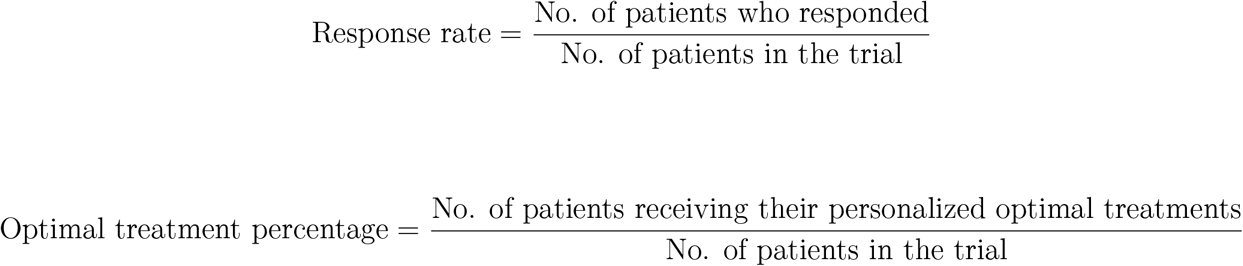

The power and the average treatment effect (ATE) adjusted by the IPTW method were also reported to thoroughly evaluate each methods’ performance. The power of a clinical trial refers to the probability of detecting a difference between different treatment groups when it truly exists; ATE was defined in the previous section (2.5). Additionally, we proposed a new criteria, the individual loss to quantify the loss for each patient due to receiving suboptimal treatments. For the individual loss, we first define a match for the enrolled patients. A match occurs when the patient’s actual treatment received is the same as the best treatment from the true model. For an enrolled patient *i* with signature *x*, let 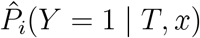 denote the probability of responding to the received treatment *T*. Let 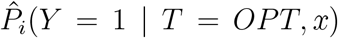 denote the probability of responding to the optimal treatment determined by the true model. Then we define the personalized loss function as follows:

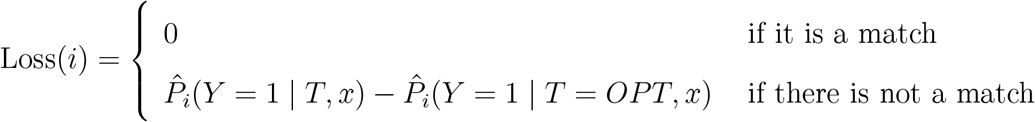

A low individual loss value suggests that the majority of patients have received the treatment and will respond at least as well as the real model’s optimal therapy.

## 3 RESULTS

### 3.1 Simulation

We used simulation studies to evaluate the proposed methods.

#### 3.1.1 Setting

We generated the *i*^*th*^ patient’s response from a logistic regression model with 10 biomarkers:

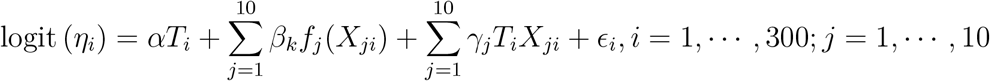

where *T*_*i*_ is the treatment indicator (either treatment 0 or treatment 1), *α* is the treatment main effect coefficient, *X*_*ji*_ is the *j*^*th*^ biomarker for patient *i*, and *f*_*j*_(*X*_*ji*_) incorporates some polynomial and step function terms. *γ*_*j*_ is the biomarker-treatment interaction coefficient and *X*_5_ ∼ *X*_6_ were assumed to interact with the treatment. A random noise for each subject is denoted as *ϵ*_*i*_. In detail, each biomarker *X*_*ji*_ was assumed to follow a normal distribution with a mean of 0 and a standard deviation of 1. Among these 10 biomarkers, *X*_2_ ∼ *X*_4_ contributed to the true model as third-degree polynomials, while *X*_7_ ∼ *X*_10_ contributed to the true model as step functions:

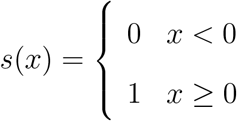

Seven scenarios of different treatment main effects (*α*=0, 0.5, 0.7, 1, 1.3, 1.5, 1.7) and a fixed treatment-biomarker interaction (*γ* = 0.5) were considered. We conducted 1000 Monte Carlo simulations for each scenario and compared the results obtained by the ML-based and ML-ensemble RAR designs with the results from the equal randomization design.

#### 3.1.2 Response rate and optimal treatment percentage

The response rate results and the percentage of receiving the optimal treatment are shown in Figure 2. Overall, the performance of ML-based RAR designs is better than the performance of the equal randomization (ER) design. When the treatment main effect is zero, the differences for both response rate and the optimal treatment percentage between ML-based RAR designs and the ER design are not significant. As the treatment effect increases, these differences become more obvious. Among these nine ML algorithms, the neural network has the highest response rate and the highest proportion of patients receiving their optimal treatments. Additionally, the ensemble method combining these nine ML methods outperforms all other methods and achieves an approximate 5% higher response rate and a more than 20% larger optimal treatment percentage compared to the ER design.

**Figure 2:**
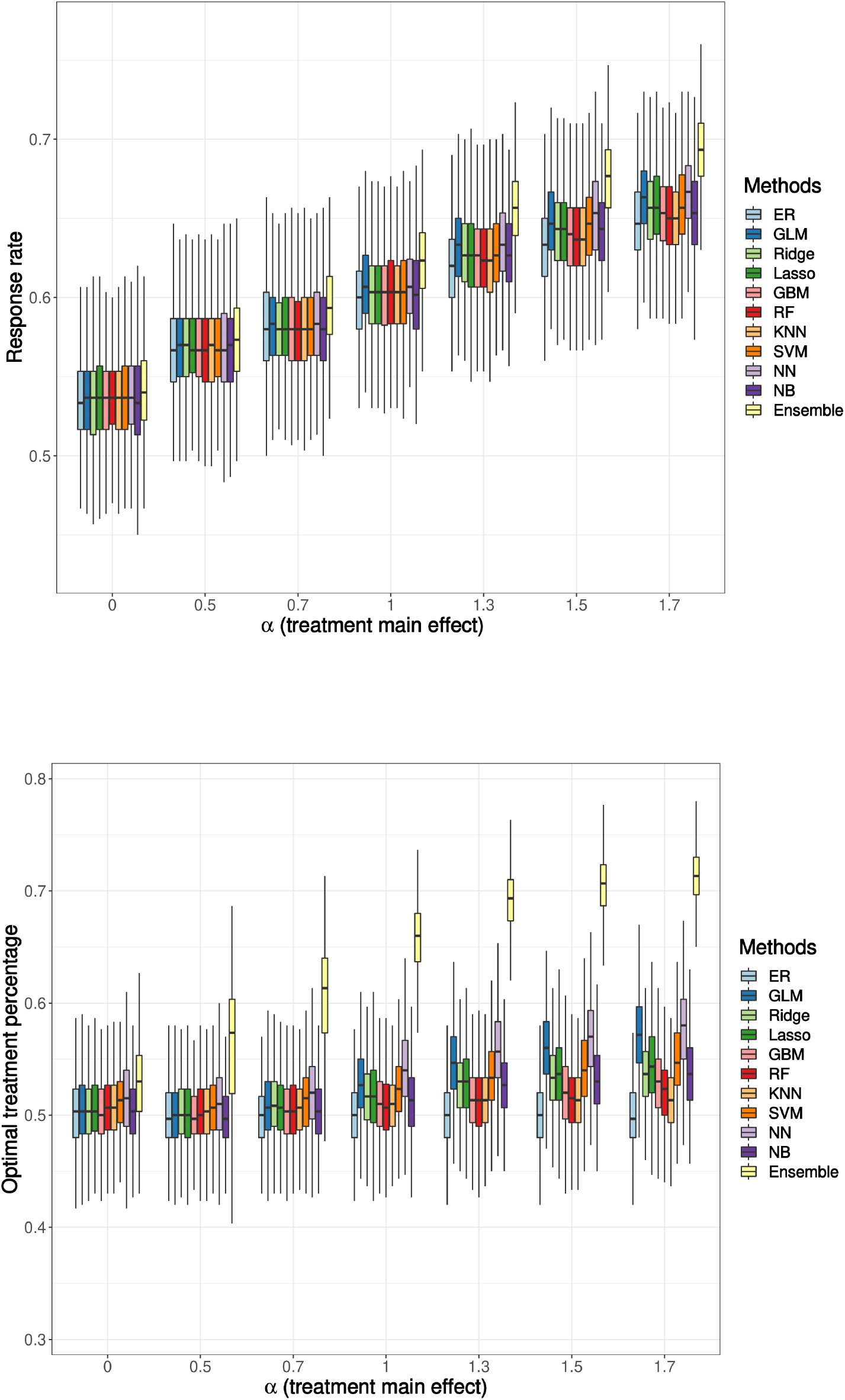
Simulation result: response rate (top), percentage of patients receiving their optimal treatments (bottom). The treatment-biomarker interaction, *γ* is fixed at 0.5. Boxplots display the median (middle line), the inter-quartile range (hinges), and 1.5 times the inter-quartile range (lower and upper whiskers) based on 1000 times simulation. The mean (over 1000 simulations) response rate ranges from 0.53 to 0.69, and the mean of optimal treatment percentages ranges from 0.50 to 0.71.

#### 3.1.3 Individual loss, ATE and power

The individual loss and the ATE results are shown in Figure 3. The interpretation of the individual loss results coincides with the previous response rate results and the optimal treatment percentage results such that the ML-ensemble RAR design has the lowest individual loss value among all scenarios, which is preferred in the trial. The ATE has been adjusted by the IPTW method to account for confounding effect of using observational data. The logistic regression method now has the highest ATE, followed by the NN method. The ensemble method has a relatively low ATE, but it is higher than the ER method when the treatment main effect becomes larger. This shows that the average effect of changing the entire population from untreated to treated using RAR designs is better than that of using the ER design.

**Figure 3:**
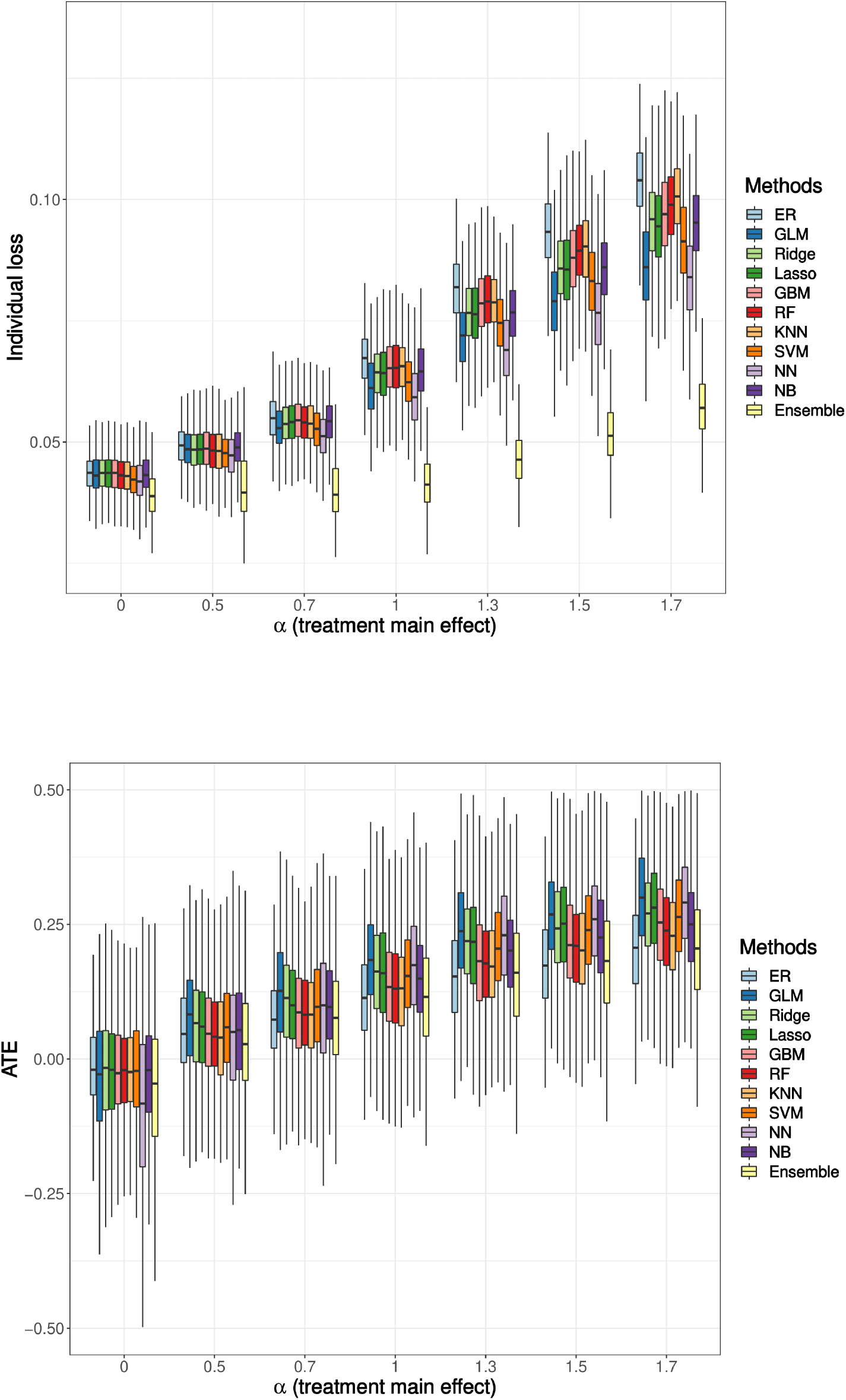
Simulation result: individual loss (top), average treatment effect (ATE, bottom). The treatment-biomarker interaction, *γ* is fixed at 0.5. Boxplots display the median (middle line), the inter-quartile range (hinges), and 1.5 times the inter-quartile range (lower and upper whiskers) based on 1000 times simulation. The mean (over 1000 simulations) individual loss ranges from 0.04 to 0.10, and the mean ATE ranges from −0.12 to 0.30.

#### 3.1.4 Power

The power results are shown in Figure 4. The power is also weighted by the IPTW method to address potential bias. For the power analysis, the Type I error is controlled at 0.05. Several papers have shown in their simulation studies that the correlation among treatment assignments was inevitable when performing inference on the data from RAR design-implemented studies [36, 44]. This correlation can increase the binomial variability and lower the power. In our simulation, the RAR design using the NN method has the lowest power, followed by using the logistic regression method. However, other ML-based RAR designs have comparable or even higher power than that of the ER design. The ensemble method has a relatively low power, but it is still better than the NN method.

**Figure 4:**
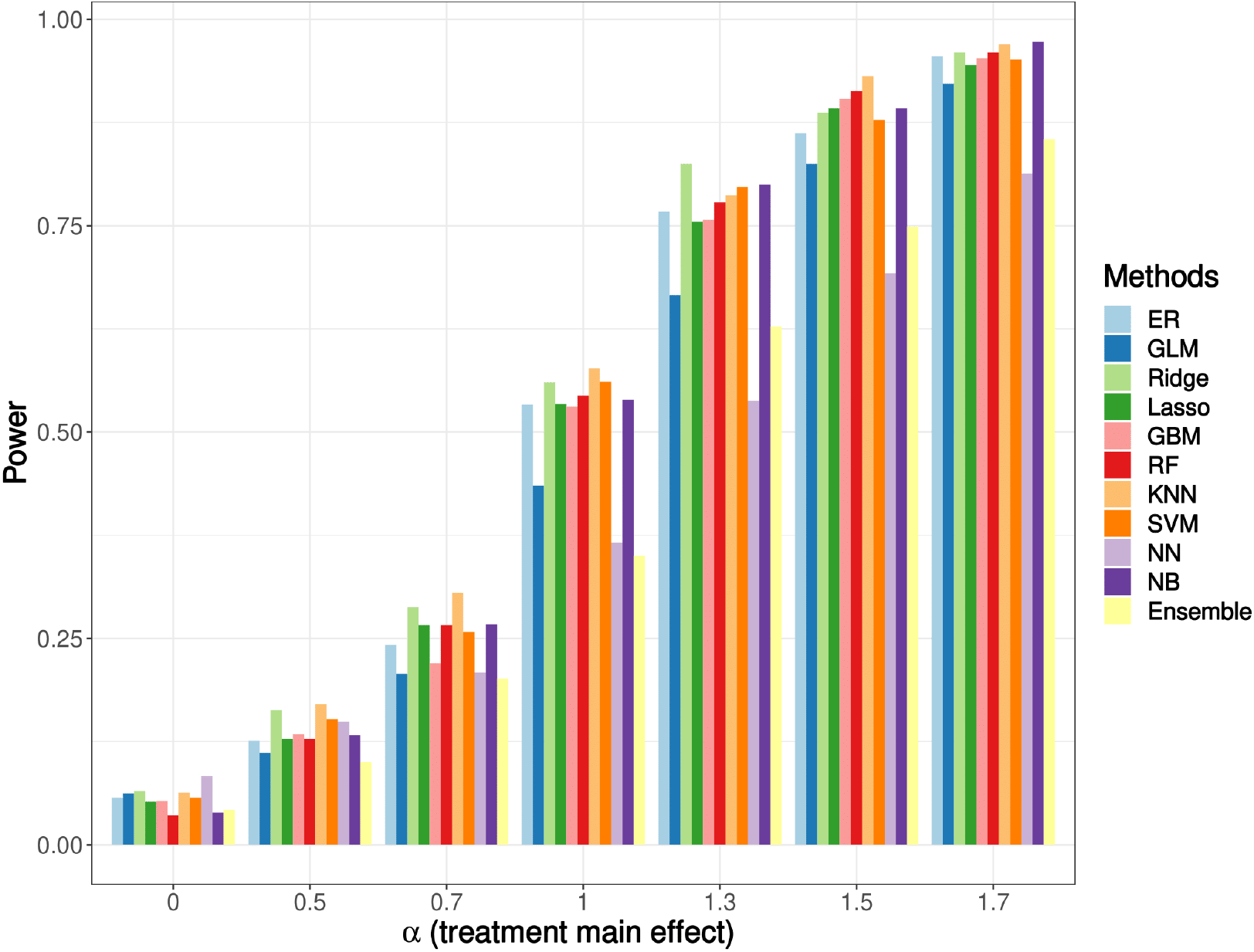
Simulation result: power. The treatment-biomarker interaction, *γ* is fixed at 0.5. The Type I error is controlled at 0.05. The power ranges from 0.04 to 0.97.

### 3.2 Real-world example

We analyzed a publicly available acute myeloid leukemia (AML) dataset from Kornblau(2009) where most of the clinical biomarkers are expression levels of cellular proteins [27]. Kornblau et al. sequenced protein expressions in leukemia-enriched cells from 256 newly diagnosed AML patients with a primary goal of eventually establishing a proteomic-based categorization of AML. The treatment and the response variables were carefully adjusted to binary variables. Specifically, the treatments were binarized to high-dose ara-C (HDAC)–based treatments and non-HDAC treatments; the responses were binarized to complete response (CR) and non-CR.

We first performed a feature selection to decide what interaction terms should be included in the model. We used each protein-treatment interaction term to build the generalized linear model (GLM) model and reported the p-value for each interaction to assess whether it has strong correlation with the dependent variable/the treatment response. The top 10 proteins whose interaction variables have the smallest p-values were selected. We then performed a gene network analysis on the genes that code for these proteins using GeneMANIA (http://genemania.org) [55]. This analysis helps to illustrate the hidden interaction and network of the corresponding genes. Additionally, it shows other genes that have been reported to associate with the input 10 genes, using extensive existing knowledge such as protein and genetic interactions, pathways, co-expression, co-localization, and protein domain similarity. The results are presented in Figure 5. The top 10 genes corresponding to the biomarkers identified in our study are highlighted with red circles.

**Figure 5:**
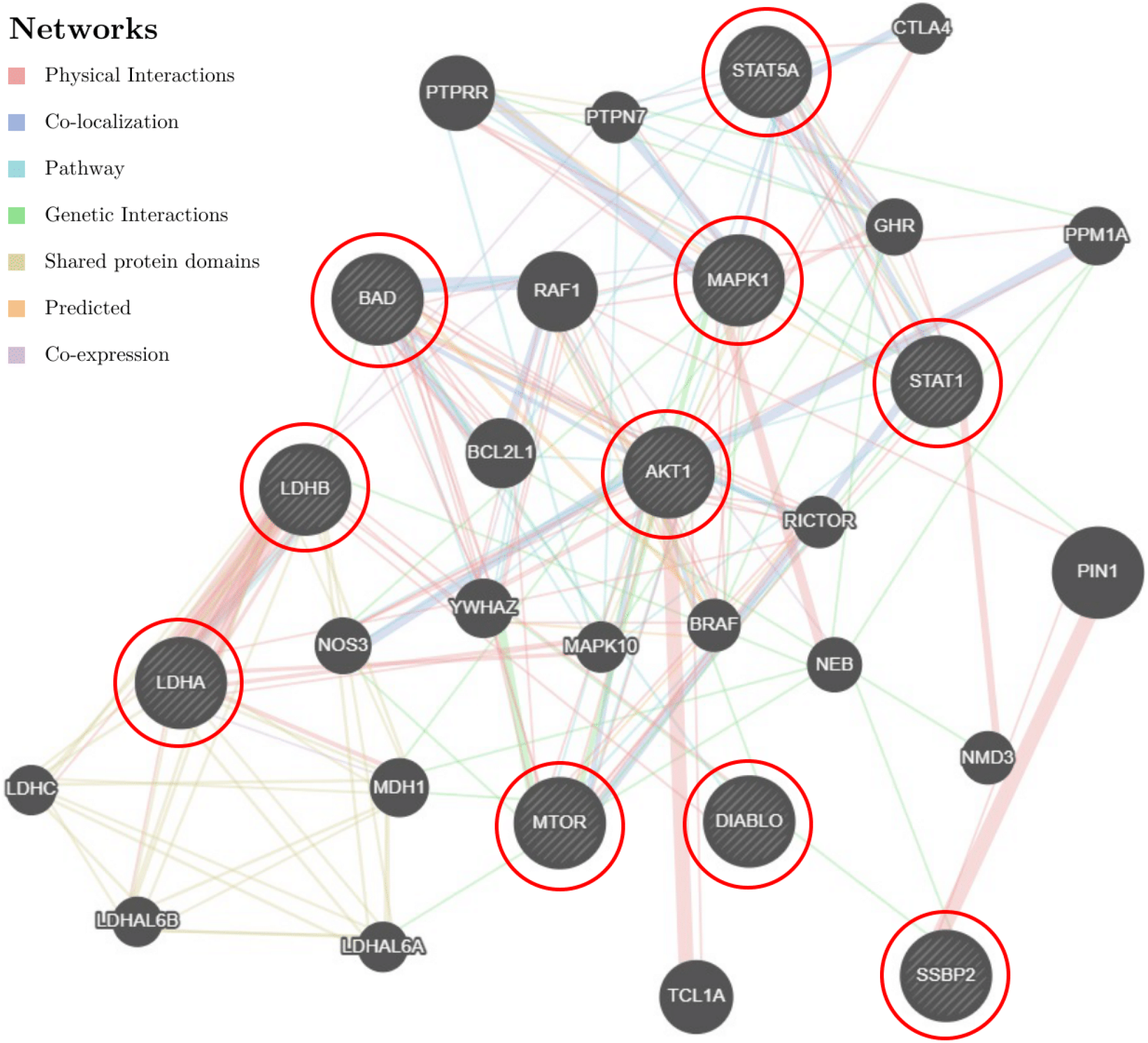
AML data: the gene network analysis. The input 10 genes, namely the genes coding for top 10 proteins that significantly interacted with the treatment, were highlighted using red circles. Other genes that were presumably involved in AML were returned by GeneMANIA.

Using a cut-off p-value of 0.1 among 71 proteins, the expression levels of three of them were found to have the most significant interactions with the treatment, i.e., the strongest correlation with the treatment outcomes: phosphothreonine 308 of Akt (Thr 308 p-Akt), the mechanistic target of rapamycin (mTOR), and signal transducer and activator of transcription 1 (STAT1). Studies have shown that these three proteins play critical roles in human AML. The level of Thr 308 p-Akt is associated with high-risk cytogenetics and predicts poor overall survival for AML patients [17]. In AML, the mTOR signaling pathway is deregulated and activated as a consequence of genetic and cytogenetic abnormalities. The mTOR inhibitors are often used to target aberrant mTOR activation and signaling [14, 45]. The STAT1 transcription factor is constitutively activated in human AML cell lines and might contribute to the autonomous proliferation of AML blasts. The inhibition of this pathway can be of great interest for AML treatments [20, 56]. Hence, we chose these three proteins to build ML models in our proposal.

The whole dataset (256 observations) was randomly shuffled and divided into two equal-sized blocks: block 1 and block 2. Each block was taken in turn as either the training set or the testing set. The results were aggregated after 100 repetitions. Since this clinical trial is already completed and it is not possible to get actual treatment responses using our methods, we separated the enrolled patients into two groups: a consistent group whose real treatments are the same as the treatments using the ML-based RAR designs and an inconsistent group whose real treatments are different from the treatments using the ML-based RAR designs. We compared the response rates in these two groups to elucidate the potential gain if the proposed RAR had been implemented. The results of each method are shown in Figure 6. In the consistent group, the response patient percentages are at least 10% higher than 50%; while the response patient percentages in the inconsistent group are all lower than 50%, i.e., we observe higher response rates in the consistent group. This means that patients in the inconsistent group may likely benefit from the RAR method we developed.

**Figure 6:**
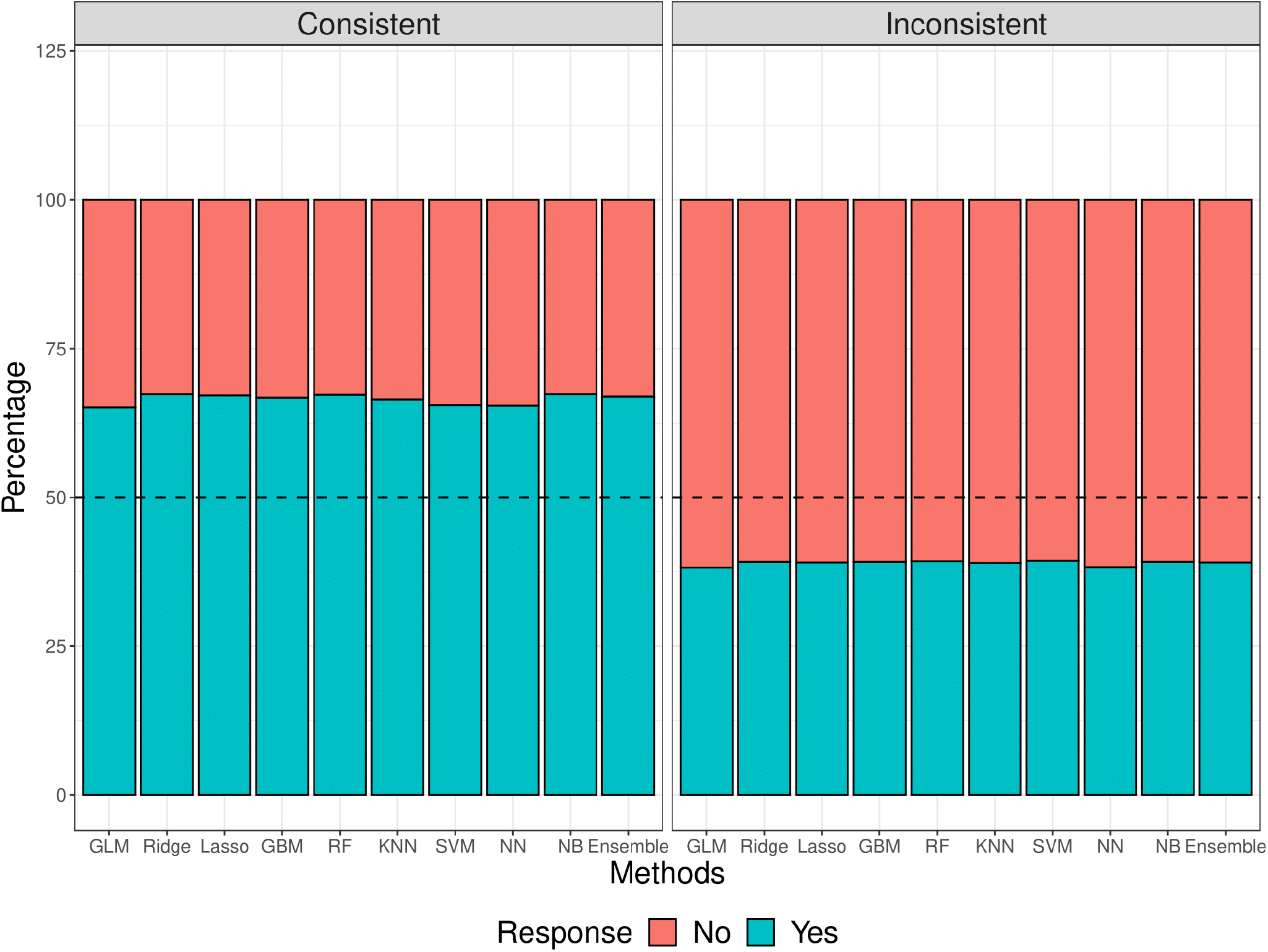
AML data result: the response percentage. Patients in the consistent group (left) were assigned to the same treatments using our ML-based RAR designs, while patients in the inconsistent group (right) were assigned to different treatments using our ML-based RAR designs. The 50% response percentage is marked with a black dashed line.

## 4 DISCUSSION

Patients are accrued in groups sequentially. RAR designs determine the treatment allocation for new groups of patients based on the accrued information of how previous groups of patients responded to their treatments. The number of RAR implementations, *k*, should be pre-defined. The choice of *k* may depend on the total sample size, trial length, and other logistics and practical considerations. Our simulation study used *k* = 2 for a total sample size of 300. In the real data analysis with a smaller sample size of 256 subjects, we used *k* = 1.

We developed novel methods for RAR designs by incorporating nine ML methods to predict treatment response and assign treatments accordingly. We showed that our ML-based RAR designs can effectively improve treatment response rates among patients. We further proposed an ensemble approach based on the consensus of the nine-ML methods to improve the prediction and decision making. Our proposed ML-ensemble RAR design builds on the predictive ability of nine ML methods and can further improve predication accuracy and patient outcome. Specifically, suppose *m* out of 9 models indicate that treatment *A* is better than treatment *B* for patient *i*, then we let *p*_*iA*_ = *m/*9 for 2 ≤ *m* ≤ 7, let *p*_*iA*_ = 0.85 for *m* ≥ 7, and *p*_*iA*_ = 0.15 for *m* ≤ 2. For *m* ≥ 7, we keep the assignment probability as a constant of 0.85 because we still want to reserve some randomness in the trial. These settings can be tuned based on prior knowledge of the treatment selections.

While we only considered settings of two treatment options in this work, ML-based RAR design can extend to multiple targeted treatments. Given *L* treatments, the *l*^*th*^ treatment allocating probability of patient *i* is shown as 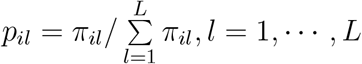, where *π*_*il*_ denotes the response probability of *l*^*th*^ treatment for patient *i* predicted by the ML algorithm. For example, NN can naturally adapt to a multi-class classification problem by replacing the binary cross-entropy loss to a categorical cross-entropy loss [19].

## 5 Conclusion

Machine learning methods have successfully demonstrated their superior prediction performance in many applications, but have not been applied to conduct RAR in clinical trials. In this study, we developed novel methods for RAR designs by incorporating ML algorithms to predict treatment response and assign treatments accordingly. We showed that the ML-based RAR designs have better performance than that of the traditional ER design. And the ensemble approach demonstrated better results than the ER design at the greatest extent. As the ML field is getting mature and abundant packages are available on different programming software, our method is easy to implement in current clinical trial systems.

## 6 Data availability

The AML dataset used for real-world illustration was downloaded from https://bioinformatics.mdanderson.org/public-datasets/supplements/ under “RPPA Data in AML” [27].

## 7 Funding

The research of Huang was partially supported by the US National Institutes of Health grants U54CA096300, U01CA253911, and 5P50CA100632, and the Dr. Mien-Chie Hung and Mrs. Kinglan Hung Endowed Professorship.

## 8 CONFLICT OF INTEREST STATEMENT

The authors have no competing interests to declare.

## Notes

### Competing Interest Statement

The authors have declared no competing interest.

